# Osteopontin Upregulation Defines a Pre-Rupture State in Thoracic Aortic Aneurysms in Mice and Humans

**DOI:** 10.64898/2026.05.27.728313

**Authors:** Kaori Sugiyama, Yuya Sato, Hiroko Matsunaga, Kenichi Kimura, Kosuke Kataoka, Toru Asahi, Hiromi Yanagisawa, Haruko Takeyama

## Abstract

**Background:** Thoracic aortic aneurysm (TAA) is a life-threatening condition with an unpredictable lisk of rupture. Current clinical parameters have limited ability to accurately predict imminent rupture. Osteopontin (OPN) has been implicated in aortic aneurysm pathology, however, it role as a marker of imminent rupture remains. unclear. We investigated the dynamics of OPN expression dynamics in a mouse model with predictable rupture timing and validated our findings in human TAA.

**Methods:** One-month-old fibrillin-1 hypomorphic (*Fbn1*^mgR/mgR^) mice were used as a TAA model; with wild-type (WT) mice served as controls. Angiotensin II (AngII) was administered to *Fbn1*^mgR/mgR^ to induce acute aortic rupture. Single-section transcriptome analysis and immunofluorescence staining were performed on ascending aortic tissue at 24 and 72 hours after AngII infusion, with pre-treatment *Fbn1*^mgR/mgR^ and WT mice serving as controls. To determine conservation in human disease, we reanalyzed publicly available single-cell RNA sequencing data from ascending thoracic aortic aneurysm (ATAA) patients.

**Results:** AngII infusion induced progressive mortality beginning at 24 hours, with approximately 60% survival at 72 hours and nearly no survival by 8 days in *Fbn1^mgR/mgR^* mice. At this pre-rupture time point, OPN showed prominent upregulation at both mRNA and protein levels in ascending aortic tissues compared to controls. Immunofluorescence staining revealed increased OPN expression in the aortic wall, particularly in regions exhibiting structural deterioration. Reanalysis of human ATAA single-cell data showed elevated OPN expression compared to controls, with enrichment in immune cell populations, especially macrophages. Within the macrophage compartment, subcluster analysis identified a stress-responsive subpopulation (MC1) that was markedly expanded and almost exclusively composed of ATAA-derived cells, representing the primary source of OPN upregulation.

**Conclusions:** OPN upregulation represents a conserved molecular signature of the pre-rupture state in TAA across mice and humans. Our mode, which enables predictable rupture timing, allowed the capture of acute pre-rupture molecular changes, suggesting OPN as a potential biomarker for predicting imminent aortic rupture.

## Introduction

Thoracic aortic aneurysm (TAA) is a life-threatening condition that can result in sudden aortic rupture or dissection with high mortality ^1^ .Current clinical decision-making for surgical intervention relies primarily on anatomical parameters such as maximal aortic diameter and growth rate^2^. However, these morphological criteria have limited predictive value for rupture risk, as rupture can occur in relatively small aneurysms, whereas some large aneurysms remain stable for extended periods ^3^. These observations indicate that aortic rupture is not solely determined by vessel size but is strongly influenced by the molecular and cellular state of the aortic wall.

Inflammation, extracellular matrix (ECM) remodeling, and immune cell infiltration have been implicated in the pathogenesis of TAA^4^. However, most molecular studies have analyzed tissues obtained after rupture or at advanced disease stages, where secondary inflammation and tissue necrosis may obscure molecular events that directly precede rupture. Furthermore, existing transcriptomic approaches in aortic research have relied primarily on bulk tissue homogenates ^5^, spatially resolved microregion sampling^6^, or single-cell dissociation-based methods^7,8^, which either average gene expression across structurally heterogeneous regions, profile a limited gene panel, or lose tissue-level structural context. Consequently, the molecular landscape of the aortic wall immediately before rupture remains poorly defined.

Animal models provide an essential experimental platform for investigating rupture-associated mechanisms, as planned collection of pre-rupture aortic tissue in humans is ethically and practically infeasible. Fibrillin-1 hypomorphic (*Fbn1^mgR/mgR^*) mice recapitulate key features of Marfan syndrome–associated aortopathy and are widely used as a genetic model of heritable TAA ^9^. However, because spontaneous aortic rupture in *Fbn1^mgR/mgR^* mice occurs over a broad and unpredictable time window^10^, the pre-rupture period cannot be experimentally defined, precluding systematic collection of aortic tissue immediately prior to rupture. In the present study, we established an AngII-induced acute aortic rupture model in *Fbn1^mgR/mgR^* mice to control rupture timing, and developed single-section transcriptome analysis as a novel approach for directly linking the structural and molecular states of individual aortic wall sections. Using this framework, we identified osteopontin (*Spp1*/OPN) as a notably upregulated gene during the pre-rupture period, a finding further supported by reanalysis of publicly available single-cell RNA sequencing data from patients with TAA^7^.

## Methods

### Animals

All animal experiments were approved by the Institutional Animal Care and Use Committee of the University of Tsukuba and were conducted in accordance with institutional guidelines and relevant regulations (25-398). Mice were housed at the University of Tsukuba Animal Research Center under specific pathogen-free conditions with a 12-hour (h) light/dark cycle and ad libitum access to food and water. All efforts were made to minimize animal suffering and to reduce the number of animals used. For surgical procedures, mice were anesthetized using a combination of medetomidine, midazolam, and butorphanol.

### Mice Genotyping

Genomic DNA was extracted from tail biopsies using standard protocols. Genotyping of fibrillin-1 hypomorphic (*Fbn1*^mgR/mgR^) mice and wild-type (WT) littermates was performed by polymerase chain reaction (PCR) using primer sequences provided by The Jackson Laboratory. PCR products were analyzed by agarose gel electrophoresis to confirm genotypes.

### AngII-Induced Aortic Rupture Model

One-month-old male WT and *Fbn1^mgR/mgR^* mice were subcutaneously implanted with osmotic minipumps (model 1007D; Alzet Osmotic Pumps) delivering angiotensin II (AngII; A9525, Sigma-Aldrich) at a dose of 300 ng/kg/min or saline as a control for up to 7 days. In addition, pre-treatment AngII mice of both genotypes were included as controls. Mice were monitored daily for signs of distress or sudden death.

### Tissue Collection and Processing

Mice were deeply anesthetized and perfused with PBS. The thoracic aorta was carefully dissected under a stereomicroscope. Perivascular adipose tissue was gently removed, and the ascending aorta was isolated. Tissues were embedded in Tissue-Tek O.C.T. compound (Sakura Finetek) and snap-frozen. Serial transverse cryosections were prepared at a thickness of 10 μm using a cryostat. For RNA sequencing, individual sections were collected onto Kawamoto’s film (Kawamoto film type II C, Section-lab). For immunofluorescence staining, adjacent sections were mounted onto glass slides. Sections designated for RNA analysis were processed immediately or stored at −80°C until further use.

### Histological Staining

Hematoxylin and eosin (H&E) staining was performed using standard histological procedures to assess overall tissue morphology. Elastin staining was performed to visualize elastic fiber architecture in the aortic wall using established protocols^11^. Stained sections were examined using a light microscope (Axio Imager, Carl Zeiss), and representative images were acquired under identical imaging conditions for comparison among experimental groups.

### Immunofluorescence Staining and Imaging

Frozen sections were fixed with 4% paraformaldehyde for 10 min and blocked with 5% bovine serum albumin (BSA) in 0.1 % Triton X-100 phosphate-buffered saline (PBS) for 30 min at room temperature. Sections were incubated with primary antibodies diluted in blocking solution for 4 h at room temperature: CD45 (30-F11, Cell Signaling), osteopontin (AF808, R&D Systems), and *CD68* (sc-20060, Santa Cruz). After washing, sections were incubated with appropriate species-specific secondary antibodies: Cy3, Alexa 647 (Jackson ImmunoResearch) conjugated to fluorophores, together with Hoechst 33342 for nuclear counterstaining, for 30 min at room temperature. Sections were then mounted using an antifade mounting medium (ProLong Diamond Antifade Mountant, Thermo). Fluorescence images were acquired using a fluorescence microscope and a confocal laser scanning microscope. Image acquisition settings were kept constant across experimental groups. Representative images were selected to reflect the staining patterns consistently observed within each group.

### Single-section RNA sequencing preparation from Individual Aortic Wall Sections

Individual cryosections (10 μm thick) of the ascending aortic wall mounted onto Kawamoto’s film were used for RNA preparation. From each section, a defined region of interest was excised using a 2-mm biopsy punch. The punched tissue was recovered directly from the Kawamoto’s film and immediately processed to minimize RNA degradation. Because of the limited amount of starting material obtained from a single section, total RNA was not isolated or quantified. Instead, punched tissue fragments were processed using the SMART-seq2 low-input RNA preparation workflow suitable for minute tissue samples and directly subjected to downstream library preparation.

### RNA Sequencing and Library Preparation

RNA sequencing library preparation and sequencing were performed using a low-input micro-tissue RNA sequencing protocol as previously described^12^. Briefly, tissue fragments excised from individual cryosections and recovered from Kawamoto’s film were directly subjected to cDNA synthesis and library construction according to the established protocol, which minimizes sample loss during processing. Sequencing libraries were prepared following the published procedure and subjected to next-generation sequencing.

### RNA Sequencing Data Processing and Normalization

Raw sequencing reads were processed using the nf-core/rnaseq pipeline^13^ (v3.16.1). Reads were aligned to the mouse reference genome (GRCm38/mm10) using STAR aligner (v2.7.11b) ^14^. Gene-level read counts were quantified using featureCounts ^15^ (v2.0.6) with GENCODE ^16^ mouse gene annotations (vM25). Raw counts were normalized using the DESeq2 ^17^ median-of-ratios method to account for differences in library size and RNA composition.

### Differential Expression Analysis

Differential gene expression analysis was performed using DESeq2 (v1.46.0). Genes were considered differentially expressed if they met the following criteria: Benjamini-Hochberg ^18^ adjusted p-value less than 0.05 and absolute log2 fold change greater than 1. Nine pairwise comparisons were performed, including pre-treatment versus AngII 24 h within each genotype, pre-treatment versus AngII 72 h within each genotype, AngII 24 h versus AngII 72 h within each genotype, and WT versus *Fbn1^mgR/mgR^*at AngII pre-treatment, 24 h, and 72 h time points. UpSet plots were generated using the ComplexUpset R package (v1.3.3) ^19^ to visualize the intersection patterns of DEGs across multiple comparisons, with the top 12 intersections ranked by set size displayed.

### Time-Series Clustering Analysis

Soft clustering of temporal gene expression profiles was performed using the Mfuzz R package (v2.70.0)^20^. The analysis was conducted separately for WT and *Fbn1^mgR/mgR^* mice samples. Prior to clustering, outlier data points were removed using an interquartile range (IQR)-based filtering method. For each gene at each time point within each genotype, values exceeding Q1 minus 1.5 times IQR or Q3 plus 1.5 times IQR were excluded, where Q1 and Q3 represent the first and third quartiles, respectively. Genes with standard deviation less than 1.5 across time points were excluded prior to clustering to focus on dynamically expressed genes. Expression values were then averaged across biological replicates for each time point and standardized using z-score transformation prior to clustering. The number of clusters was set to 7. The fuzzifier parameter was automatically estimated using the mestimate function. Genes were assigned to clusters based on their maximum membership score, with a minimum threshold of 0.25 for inclusion in downstream analyses.

### Cluster Overlap Analysis

The statistical significance of gene overlap between WT and *Fbn1^mgR/mgR^* mice clusters was assessed using the hypergeometric test (one-tailed). For each pair of clusters, the test evaluates whether the observed number of shared genes exceeds what would be expected by chance. The p-value was calculated using the phyper function in R, where k is the observed overlap, m is the size of the first cluster, N is the total number of unique genes (N = 33,117), and n is the size of the second cluster. Results were visualized as -log10(p-value) in a heatmap using ggplot2.

### Gene Ontology Enrichment Analysis

GO enrichment analysis^21^ was performed using the clusterProfiler R package (v4.18.4) ^22^ with the org.Mm.eg.db annotation database (v3.22.0). Gene symbols were converted to Entrez IDs using the bitr function. *Fbn1^mgR/mgR^*mice-specific genes were defined as genes present in a given *Fbn1^mgR/mgR^* cluster but absent from the corresponding WT mice cluster based on the cluster pairing defined by the overlap analysis. Enrichment analysis was performed for Biological Process ontology with a p-value cutoff of 0.05 and q-value (FDR) cutoff of 0.2 using the Benjamini-Hochberg method for multiple testing correction. Minimum and maximum gene set sizes were set to 10 and 500, respectively.

### Differential Response Gene Identification

To identify genes with the most pronounced genotype-specific temporal responses, we calculated the difference in mean expression between *Fbn1^mgR/mgR^*and WT mice at the 72 h time point for all genes within each cluster. The difference score was defined as mean expression in *Fbn1^mgR/mgR^* mice at 72 h minus mean expression in WT mice at 72 h. Genes were ranked by this difference score in descending order. The top 6 genes with the largest positive difference, indicating *Fbn1^mgR/mgR^* mice-specific upregulation, were selected for detailed visualization. Temporal expression profiles were plotted using ggplot2 with the ggsci color palette.

### Human Tissue Samples and Single-Cell RNA Sequencing Data

Single-cell RNA sequencing data from human thoracic aortic tissue was obtained from the Gene Expression Omnibus (GEO) database under accession number GSE155468. The dataset comprised 11 samples, including 3 Control samples (GSM4704931, GSM4704932, GSM4704933) and 8 ATAA samples (GSM4704934, GSM4704935, GSM4704936, GSM4704937, GSM4704938, GSM4704939, GSM4704940, GSM4704941). Raw count matrices were processed as described below.

### Data Preprocessing and Quality Control

Raw count matrices were converted to AnnData format using Scanpy (v1.11.5) ^23^. Quality control filtering was applied to remove low-quality cells and genes. Genes expressed in fewer than 10 cells were excluded from the analysis. The data was normalized to 10,000 counts per cell using the total-count normalization method, followed by log1p transformation. Highly variable genes were identified using the Scanpy highly_variable_genes function with batch-aware selection across samples, selecting the top 2,000 variable genes for downstream analysis.

### Dimensionality Reduction and Batch Correction

Principal component analysis (PCA) was performed on the log-normalized expression matrix using 50 principal components. Batch effects arising from individual samples were corrected using Harmony integration (harmonypy v0.2.0)^24^ with the sample identifier as the batch key. Following batch correction, a k-nearest neighbor graph was constructed using 15 neighbors and 40 principal components from the Harmony-corrected embedding. UMAP ^25^ coordinates were computed for visualization purposes.

### Cell Type Annotation

Cell types were predicted using a weighted marker gene scoring approach. A curated panel of marker genes with positive and negative direction weights was used for eight major cell types, including Vascular Smooth Muscle Cells (positive markers: *ACTA2, MYH11, CNN1, TAGLN, MYLK*), Fibroblasts (*COL1A1, COL1A2, PDGFRA, DCN, LUM*), Endothelial Cells (*PECAM1, VWF, CDH5, KDR, VCAM1*), Macrophages (*CD68, CD14, CD163, CSF1R, AIF1*), T Cells (*CD3D, CD3E, CD8A, CD4, IL7R*), B Cells (*CD79A, CD79B, MS4A1, IGHM, CD19*), NK Cells (*KLRC1, NCAM1, KLRB1, NKG7, GZMA*), and Dendritic Cells (*ITGAX, CD1C, CLEC9A, FCER1A, IRF8*). For each cell type, a weighted score was calculated by summing the expression levels of marker genes multiplied by their respective weights, with negative markers subtracted from the score. Cells were assigned to the cell type with the highest weighted score above a minimum threshold of 0.08, otherwise classified as Unknown.

### Macrophage Subcluster Analysis

Macrophages identified from the cell type annotation were extracted for subcluster analysis. A new k-nearest neighbor graph was constructed using 15 neighbors and 20 principal components. Leiden clustering (leidenalg v0.11.0) ^26^ was performed at resolution 0.5, yielding 8 subclusters (MC1-MC8). UMAP embedding was recomputed specifically for the macrophage population.

### *SPP1* Expression Analysis

Differential gene expression between ATAA and Control conditions was assessed at both the cell level and pseudobulk level. Cell-level comparisons were performed using the Wilcoxon rank-sum test implemented in Scanpy. For *SPP1* expression analysis, log-normalized expression values were compared between conditions using the Wilcoxon test, with p-values less than 0.05 considered statistically significant. Significance levels were denoted as follows: p < 0.05 (*), p < 0.01 (**), p < 0.001 (***), and p < 0.0001 (****).

### Data Visualization

All visualizations were generated using R (v4.5.1) with the ggplot2 package (v4.0.0). Color palettes were obtained from the ggsci package (v4.2.0), using the Nature Publishing Group (NPG) palette for condition comparisons and the D3 category10 palette for cell type and subcluster coloring. UMAP plots were generated using point-based representations with black outlines for improved visibility. Violin plots were combined with boxplots to show both the distribution and summary statistics of expression values. Pie charts were created using polar coordinate transformations with percentage labels positioned within segments for values greater than or equal to 5% and external labels with leader lines for smaller segments. Heatmaps were generated using ggplot2 with a diverging color scale (blue-white-red) centered at zero. Statistical comparisons on violin plots were added using the ggpubr package with significance brackets.

### Software and Statistical Analysis

All computational analyses were performed in R (v4.5.1). The following R packages were used: DESeq2 (v1.46.0) for differential expression analysis, Mfuzz (v2.70.0) for soft clustering, clusterProfiler (v4.18.4) for GO enrichment analysis, ComplexUpset (v1.3.3) for UpSet plot visualization, ggplot2 (v4.0.0) for data visualization, cowplot (v1.2.0) for plot arrangement, and tidyverse (v2.0.0) for data manipulation. For expression profile plots, error bars represent standard deviation calculated from biological replicates after IQR-based outlier removal. Statistical significance thresholds were set at p < 0.05 unless otherwise specified.

## Results

### Establishment of an AngII-induced acute aortic rupture model in *Fbn1^mgR/mgR^ mice*

To establish an experimental model of acute aortic rupture, fibrillin-1 hypomorphic (*Fbn1*^mgR/mgR^) mice were infused with angiotensin II (AngII) or not (pre-treatment) and monitored for survival, with wild-type (WT) mice included for comparison. Kaplan–Meier survival analysis demonstrated that AngII administration induced rapid and reproducible mortality specifically in *Fbn1^mgR/mgR^*mice (n = 16), whereas pre-treatment *Fbn1^mgR/mgR^* mice (n = 6) and both pre-treatment WT mice (n = 9) and AngII-treated WT mice (n = 10) exhibited no mortality during the observation period (Figure 1A). Notably, all AngII-treated *Fbn1^mgR/mgR^*mice survived for the first 24 h following AngII administration, after which survival progressively declined. Approximately 60% of AngII-treated *Fbn1^mgR/mgR^*mice remained alive at 72 h, defining a reproducible period preceding fatal rupture. All AngII-treated *Fbn1^mgR/mgR^* mice died within 8 days of treatment, and survival differed significantly between AngII-treated *Fbn1^mgR/mgR^*mice and all pre-treatment groups (log-rank test).

**Figure 1.**
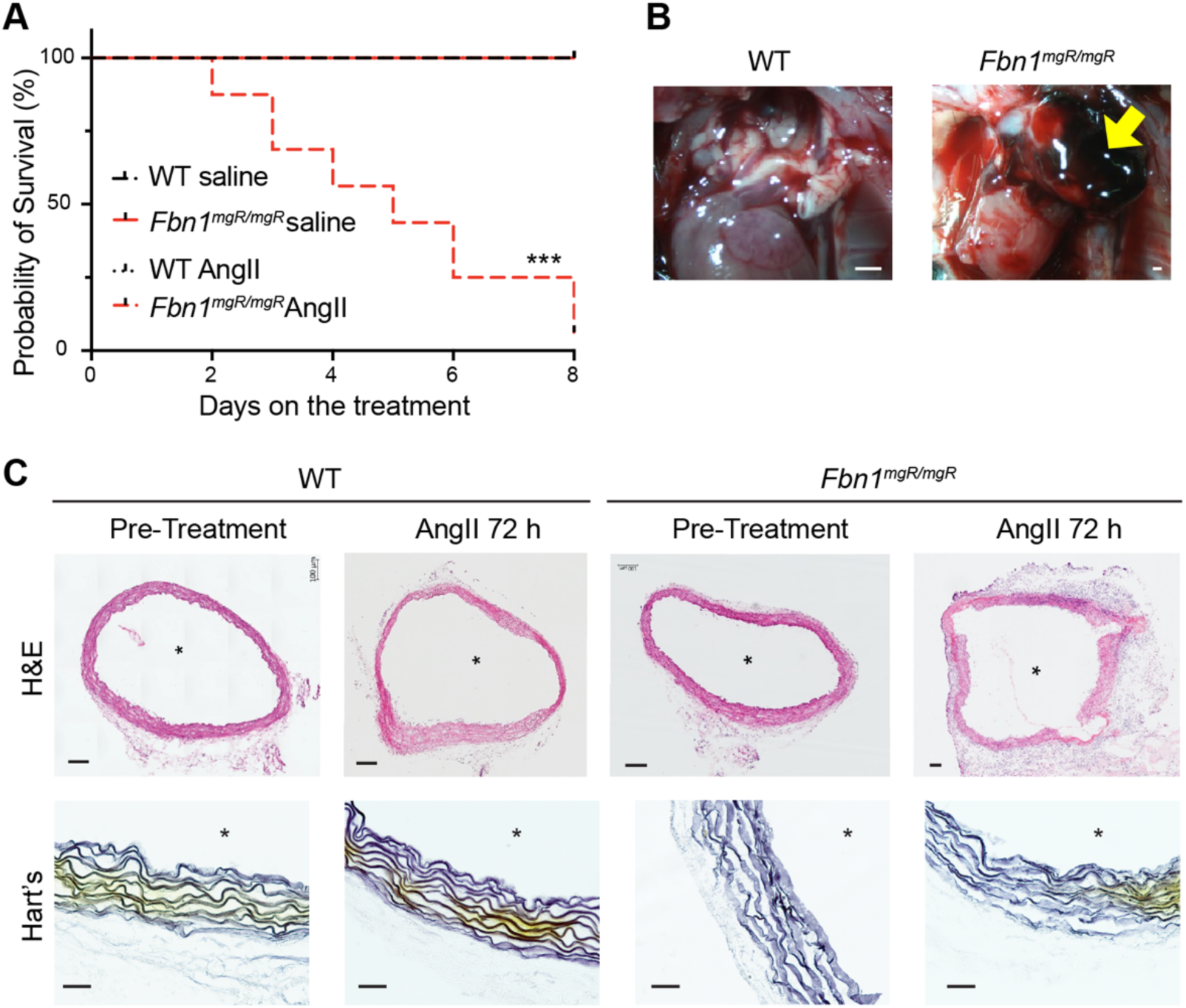
Controllable aortic rupture model induced by Angiotensin II in *Fbn1^mgR/mgR^*. A. Kaplan-Meier survival curve of WT mice (saline, n=9; Angiotensin II (AngII), n=10) and *Fbn1mgR/mgR* (saline, n=6; AngII, n=16) administered with or without AngII beginning at 1 month of age for 1 week. B. Representative gross morphology of WT and *Fbn1^mgR/mgR^* following AngII treatment. *Fbn1mgR/mgR* showed aortic rupture and/or dissection. Scale bars are equal to 1 mm. C. Histological images of aortic tissues of WT and *Fbn1^mgR/mgR^*. Hematoxylin and eosin (H&E) staining and elastin staining (Hart’s staining) for pre-treatment and AngII 72 h of WT and *Fbn1^mgR/mgR^*. Scale bars equal to 100 µm for H&E and 20 µm for Hart’s staining. *; lumen side.

Representative gross images of a WT mouse and an AngII-treated *Fbn1^mgR/mgR^* mice showed aortic rupture acute disruption of the thoracic aorta, accompanied by extensive periaortic hemorrhage (Figure 1B). Rupture sites were predominantly localized to the ascending thoracic aorta. We therefore selected 72 h post-AngII infusion as the tissue collection timepoint, as it represents a window in which the majority of mice remained alive, yet rupture was clearly imminent.

Histological analyses further demonstrated severe structural deterioration of the aortic wall following AngII treatment. Hematoxylin and eosin (H&E) staining revealed marked medial degeneration and aortic wall disruption in AngII-treated *Fbn1^mgR/mgR^* mice at 72 h post-AngII infusion, whereas pre-treatment *Fbn1^mgR/mgR^* and WT mice preserved normal medial structure (Figure 1C). Consistently, Hart’s staining showed prominent fragmentation of elastic fibers in regions of medial degeneration AngII-treated *Fbn1^mgR/mgR^*mice, while elastic fiber organization remained intact in control groups. Together, these findings demonstrate that AngII infusion selectively induces reproducible acute thoracic aortic rupture in *Fbn1^mgR/mgR^* mice and establishes a defined pre-rupture period suitable for subsequent molecular analyses.

### Time-resolved transcriptome analysis reveals *Spp1* as a pre-rupture transcriptional signature in *Fbn1^mgR/mgR^* mice

To capture transcriptional changes associated with localized structural alterations of the aortic wall preceding rupture, we employed a novel single-section transcriptome approach. By analyzing individual aortic wall sections rather than pooled tissue samples, this strategy enables direct comparison of transcriptional states between structurally preserved regions and regions exhibiting early structural compromise. This section-level analysis is particularly suited to interrogate gene expression changes occurring during the transition from an intact aortic wall to a partially compromised pre-rupture state. We designed a time-resolved single-section transcriptome analysis across defined stages following AngII administration. Individual aortic wall cryosections were collected from WT mice pre-treatment, AngII for 24 h, or AngII for 72 h, as well as from *Fbn1^mgR/mgR^*mice pre-treatment, AngII for 24 h, or AngII for 72 h, and subjected to RNA sequencing (Figure 2A). For mice in total, 30 mice were used across six experimental groups: WT mice with pre-treatment (n = 5 mice, 10 sections), WT mice with AngII 24 h (n = 5 mice, 10 sections), WT mice with AngII 72 h (n = 5 mice, 10 sections), *Fbn1^mgR/mgR^* mice with pre-treatment (n = 5 mice,10 sections), *Fbn1^mgR/mgR^* mice with AngII 24 h (n = 5 mice, 10 sections), and *Fbn1^mgR/mgR^* mice with AngII 72 h (n = 5 mice, 10 sections). Multiple serial sections (typically 2) were analyzed per mouse, yielding a total of 60 individual aortic sections for RNA sequencing. A total of 33,117 genes were detected across all samples. This experimental design enabled assessment of gene expression changes during the transition from a fully surviving state at 24 h to a defined pre-rupture period at 72 h, corresponding to progressive structural compromise of the aortic wall.

**Figure 2.**
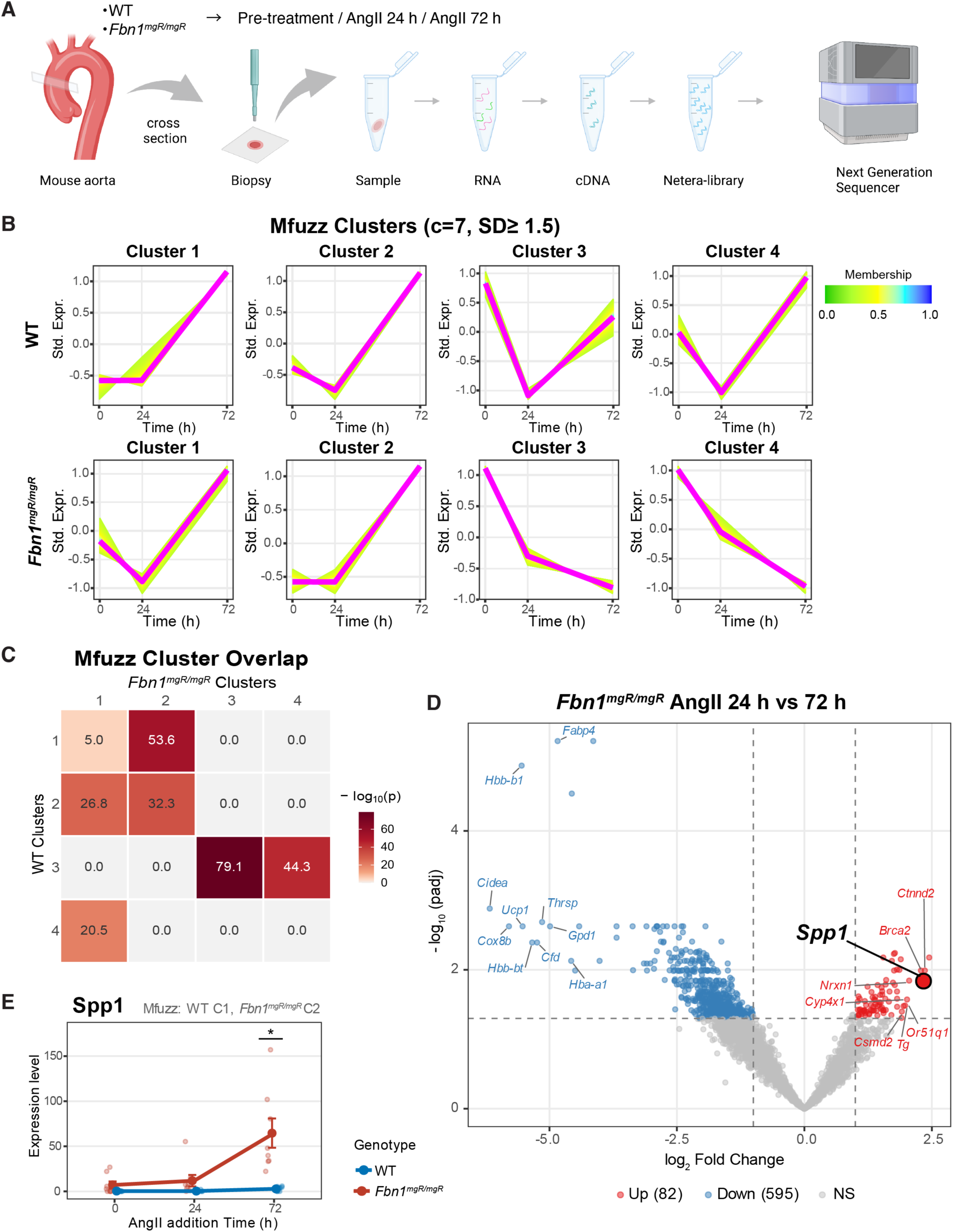
Transcriptomic analysis of individual aortic wall sections following AngII administration. A. Sample preparation for single-section RNA sequencing of mouse ascending aortic walls. RNA sequencing was performed on individual ascending aortic wall sections obtained from WT and *Fbn1^mgR/mgR^*under pre-treatment or after AngII infusion. AngII-treated samples were collected at 24 h and 72 h, corresponding to a pre-rupture stage characterized by focal aortic wall disruption without fatal rupture. B. Mfuzz soft clustering analysis of temporal gene expression patterns. Four representative clusters are shown for each genotype (WT, top row; *Fbn1*^mgR/mgR^, bottom row). Each line represents a gene, with color intensity indicating cluster membership score (0.0-1.0). The x-axis shows time after AngII administration (0, 24, 72 h), and the y-axis shows standardized expression levels (Std. Expr.). C. Heatmap showing the statistical significance of gene overlap between Mfuzz clusters of WT and *Fbn1^mgR/mgR^*. Values represent −log_10_(p-value) from hypergeometric tests. Higher values (darker red) indicate more significant overlap. Clusters 1–4 are displayed for each genotype. D. Volcano plot showing differentially expressed genes between *Fbn1^mgR/mgR^* AngII 24 h and 72 h after AngII infusion. The x-axis represents log2 fold change (*Fbn1^mgR/mgR^*AngII 24 h and 72 h), and the y-axis represents −log_10_ adjusted p-values. Red dots indicate significantly upregulated genes (82), blue dots indicate significantly downregulated genes (595), and gray dots indicate non-significant genes. *Spp1* showed marked upregulation in *Fbn1^mgR/mgR^* aortic walls. Additional upregulated genes included *Ctnnd2*, *Brca2*, *Nrxn1*, *Cyp4x1*, *Tg*, *Or51q1*, and *Csmd2*, whereas downregulated genes were *Fabp4*, *Hbb-b1*, *Cidea*, *Ucp1*, *Thrsp*, *Cox8b*, *Hbb-bt*, *Gpd1*, *Cfd*, and *Hba-a1*. Dashed vertical lines indicate log2 fold change thresholds, and the horizontal dashed line indicates the statistical significance threshold. E. Time-dependent upregulation of *Spp1* in *Fbn1^mgR/mgR^* aortic walls after AngII infusion. *Spp1* expression levels in WT and *Fbn1^mgR/mgR^* at 0, 24, and 72 h following AngII infusion. Expression values were derived from single-section transcriptomic analysis. *Spp1* belongs to WT Cluster 1 and *Fbn1^mgR/mgR^* Cluster 2 in Mfuzz analysis (Figure 2B, C). *Spp1* expression was markedly increased in *Fbn1^mgR/mgR^* at 72 h, whereas WT showed minimal changes over time. Data are presented as mean ± SEM, with individual data points shown. *p < 0.05.

Differential expression analysis using DESeq2 identified numerous genes responding to AngII stimulation in both genotypes, with distinct subsets of differentially expressed genes (DEGs) unique to each genotype and time point comparison. UpSet analysis of the intersection patterns revealed that the largest gene sets were specific to individual comparisons rather than shared across multiple conditions, with the pre-treatment vs AngII 72 h comparison in *Fbn1^mgR/mgR^*mice yielding the greatest numbers of both up-and down-regulated DEGs, consistent with maximal transcriptional change at the pre-rupture time point (Supplementary Figure S1). The number of genotype-specific DEGs increased progressively from 24 h to 72 h, suggesting a widening divergence of the transcriptional landscape between WT and *Fbn1^mgR/mgR^* mice aortas over time.

To classify temporal gene expression dynamics systematically, we applied Mfuzz soft clustering independently to WT and *Fbn1*^mgR/mgR^ mice datasets. Seven clusters were identified for each genotype, capturing diverse temporal expression patterns including upward, downward, and transient profiles (Clusters 1 – 4: Figure 2B, Clusters 5 – 7: Supplementary Figure S2A), revealing both conserved and genotype-specific responses to AngII. Cluster membership scores indicated the degree of gene association with each temporal pattern, with higher scores reflecting stronger pattern adherence.

We next examined the extent of shared transcriptional programs between genotypes by performing hypergeometric tests on all pairs of WT and *Fbn1^mgR/mgR^*mice clusters (Clusters 1 – 4: Figure 2C, all clusters: Supplementary Figure S2B). Among these clusters, WT Cluster 3 and *Fbn1^mgR/mgR^* Cluster 3 showed the strongest overlap (−log10(p) = 79.1); WT Cluster 3 exhibited a transient decrease at 24 h followed by recovery at 72 h, whereas *Fbn1^mgR/mgR^* Cluster 3 displayed a progressive downward trend. WT Cluster 1 and *Fbn1^mgR/mgR^* Cluster 2 also showed significant overlap (−log10(p) = 53.6), with both clusters displaying upward expression trends at 72 h.

Gene Ontology (GO) enrichment analysis of these overlapping clusters revealed distinct biological processes (Supplementary Figure S3). WT Cluster 3 was enriched for extracellular matrix organization and connective tissue development, whereas *Fbn1^mgR/mgR^*Cluster 3, despite sharing significant gene overlap, was instead dominated by protein stabilization and proteostasis-related pathways. WT Cluster 1 and *Fbn1^mgR/mgR^*Cluster 2 were both enriched for cilium-related processes, consistent with their concordant upward expression trends at 72 h.

To identify genes specifically upregulated during the transition to the pre-rupture state, we performed pairwise DEG analysis across all time points and genotypes (Figure 2D & Supplementary Figure S4) and focused on the *Fbn1^mgR/mgR^*AngII 24 h vs 72 h comparison. Volcano plot analysis identified 82 upregulated and 595 downregulated genes at 72 h relative to 24 h in *Fbn1^mgR/mgR^* mice. Among the upregulated genes, *Spp1* was identified as a notably upregulated gene, alongside *Brca2, Csmd2, Ctnnd2, Cyp4x1, Or51q1*, and *Tg* (Figure 2D).

Examination of expression patterns across all time points in WT and *Fbn1^mgR/mgR^* revealed that among the top upregulated genes, only Brca2 and *Spp1* showed significant differences between genotypes over the time course (Figure 2E & Supplementary Figure S5). Brca2 showed significant differences between WT and *Fbn1^mgR/mgR^* at both pre-treatment (0 h) and 72 h, suggesting a pre-existing genotypic difference rather than a rupture-specific response. In contrast, *Spp1* showed no significant difference between genotypes at pre-treatment (0 h), but was markedly and significantly upregulated in *Fbn1^mgR/mgR^* mice at 72 h following AngII administration. Among the top upregulated genes, *Spp1* was therefore the only gene demonstrating significance exclusively at the pre-rupture time point without a pre-existing baseline difference, establishing it as a candidate pre-rupture transcriptional marker in *Fbn1^mgR/mgR^* mice.

### Immunofluorescence Confirms Osteopontin Protein Upregulation at the Pre-Rupture Stage in *Fbn1^mgR/mgR^* mice

Following the identification of *Spp1* as a candidate pre-rupture transcriptional marker, we next validated its protein product, osteopontin (OPN), at the protein level by immunofluorescence staining. Aortic sections from WT and *Fbn1^mgR/mgR^* mice under pre-treatment, AngII 24 h, and AngII 72 h conditions were stained for OPN and CD45 (Figure 3). OPN expression was markedly increased exclusively in *Fbn1^mgR/mgR^* AngII 72 h, whereas all other conditions exhibited minimal OPN expression, consistent with the time-dependent mRNA upregulation observed in the transcriptome analysis. This temporal specificity at the protein level further supports OPN as a marker specifically associated with the pre-rupture state rather than a constitutive or early AngII response.

**Figure 3.**
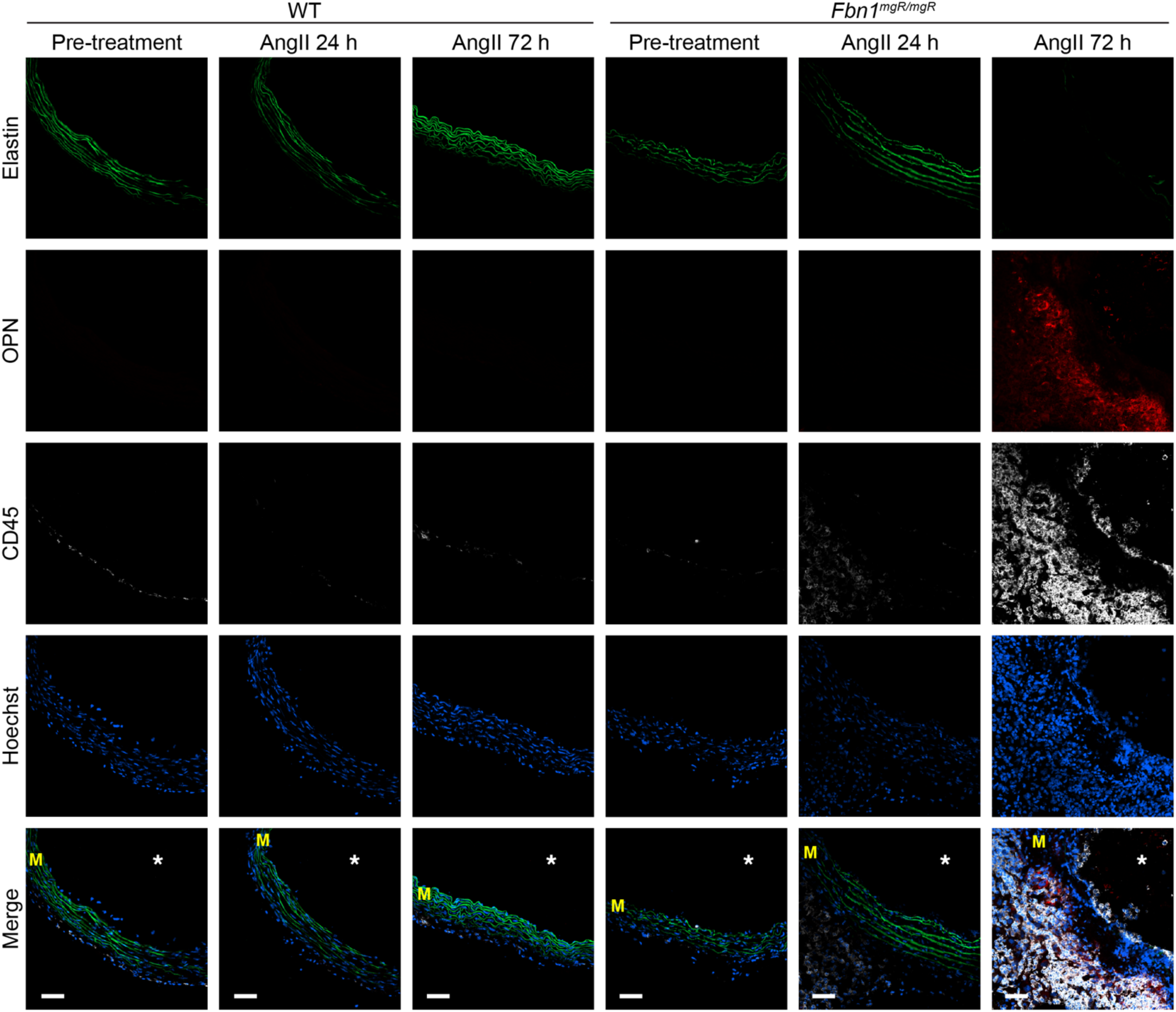
Immunofluorescence staining validation of osteopontin upregulation in the aortic wall of mouse thoracic aneurysm models. Representative immunofluorescence images of ascending aortic sections from both WT and *Fbn1^mgR/mgR^* under pre-treatment, AngII 24 h, and AngII 72 h conditions (green: elastin autofluorescence, red: osteopontin (OPN), white: CD45, blue: Hoechst). Increased OPN protein expression is observed in the aortic wall at the pre-rupture stage (AngII 72 h) in *Fbn1^mgR/mgR^*, in association with regions of structural elastin deterioration and immune cell accumulation. M, media; *, lumen. Scale bar, 25 µm.

Notably, the aortic media exhibited disrupted elastic fibers at the pre-rupture stage, while OPN expression was predominantly localized to the adventitia, with possible sparse expression near the endothelium. Co-staining with CD45, a pan-leukocyte marker expressed on immune cells including macrophages, revealed that OPN-positive regions were associated with immune cell accumulation. CD45-positive cells were also detectable in *Fbn1^mgR/mgR^*AngII 24 h, though OPN expression remained minimal at this time point, suggesting that immune cell infiltration precedes or coincides with OPN induction at the pre-rupture stage. Together, these findings confirm OPN protein upregulation at the pre-rupture stage and suggest a potential role in immune cell recruitment during aortic wall deterioration.

### OPN Expression is Upregulated in Macrophages in Human Ascending Thoracic Aortic Aneurysm

To characterize the cellular landscape of human ATAA at single-cell resolution, we reanalyzed publicly available single-cell RNA sequencing data from thoracic aortic tissue comprising 3 Control and 8 ATAA cases. After quality control and filtering, 39,869 cells were retained (Control, 7,645; ATAA, 32,224) and annotated into eight major cell populations—SMCs, fibroblasts, endothelial cells, macrophages, T cells, B cells, NK cells, and dendritic cells—whose composition differed substantially between conditions (Figure 4A & 4B). Control aortic tissue was dominated by SMCs and macrophages, reflecting the structural and resident immune composition of the normal aortic wall, whereas ATAA tissue exhibited a marked shift toward immune cell populations, with T cells, macrophages, and NK cells constituting the majority, while SMCs were dramatically reduced.

**Figure 4.**
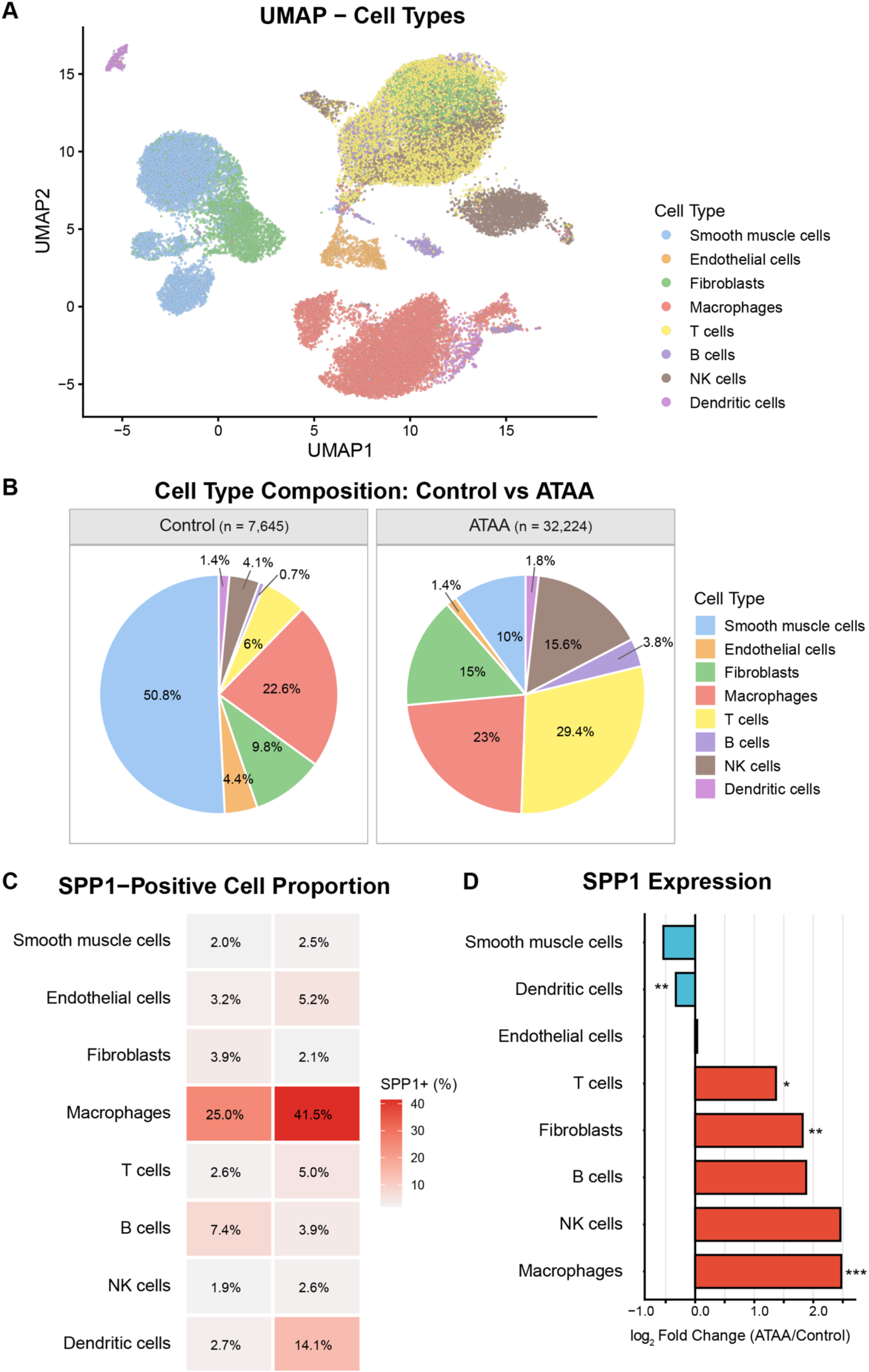
*SPP1*-expressing macrophages are expanded in human ascending thoracic aortic aneurysm. A. UMAP visualization of 39,869 cells from control (n = 7,645) and ATAA (n = 32,224) human aortic tissue, colored by annotated cell type: smooth muscle cells, endothelial cells, fibroblasts, macrophages, T cells, B cells, NK cells, and dendritic cells. B. Pie charts showing the proportional composition of cell types in Control and ATAA samples. C. Heatmap depicting the proportion of *SPP1*-positive cells within each annotated cell type in Control and ATAA. Color intensity reflects the percentage of *SPP1*-expressing cells within each population. D. Bar plot showing *SPP1* expression across annotated cell types, represented as log₂ fold change (ATAA/Control). Red and blue bars indicate upregulation and downregulation in ATAA, respectively. Statistical significance was determined by Wilcoxon rank-sum test. (\**p* < 0.05; \*\**p* < 0.01; \*\*\**p* < 0.001.)

Having identified OPN as a notably upregulated protein in *Fbn1^mgR/mgR^*at the pre-rupture timepoint, we asked whether its encoding gene *SPP1* is similarly upregulated in human ATAA. Quantification of the proportion of *SPP1*-positive cells within each annotated population revealed that macrophages harbored by far the highest *SPP1* positivity, with a marked increase in ATAA relative to Control (Figure 4C). Differential expression analysis further confirmed that *SPP1* was most significantly upregulated in macrophages among all cell types (Figure 4D), whereas SMCs and dendritic cells showed modest downregulation in ATAA.

Strikingly, although the overall proportion of macrophages was comparable between Control and ATAA,both the fraction of *SPP1*-expressing cells within the macrophage compartment and the per-cell expression level of *SPP1* increased substantially in disease, indicating a qualitative phenotypic shift within a stable macrophage population rather than simple macrophage expansion. We therefore performed subcluster analysis of the macrophage compartment to determine whether this upregulation was driven by a specific subpopulation.

### Expansion of an ATAA-Specific Stress-Responsive Macrophage Subpopulation with high OPN expression

To dissect the heterogeneity of the macrophage compartment in ATAA, we performed subclustering of 9,141 macrophages, identifying eight distinct subclusters (MC1–MC8) by Leiden clustering (Figure 5A, left panel). Visualization of *SPP1* expression on the UMAP embedding revealed that high *SPP1* expression was not uniformly distributed across macrophages but was concentrated in a discrete region corresponding to MC1 (Figure 5A, right panel).

**Figure 5.**
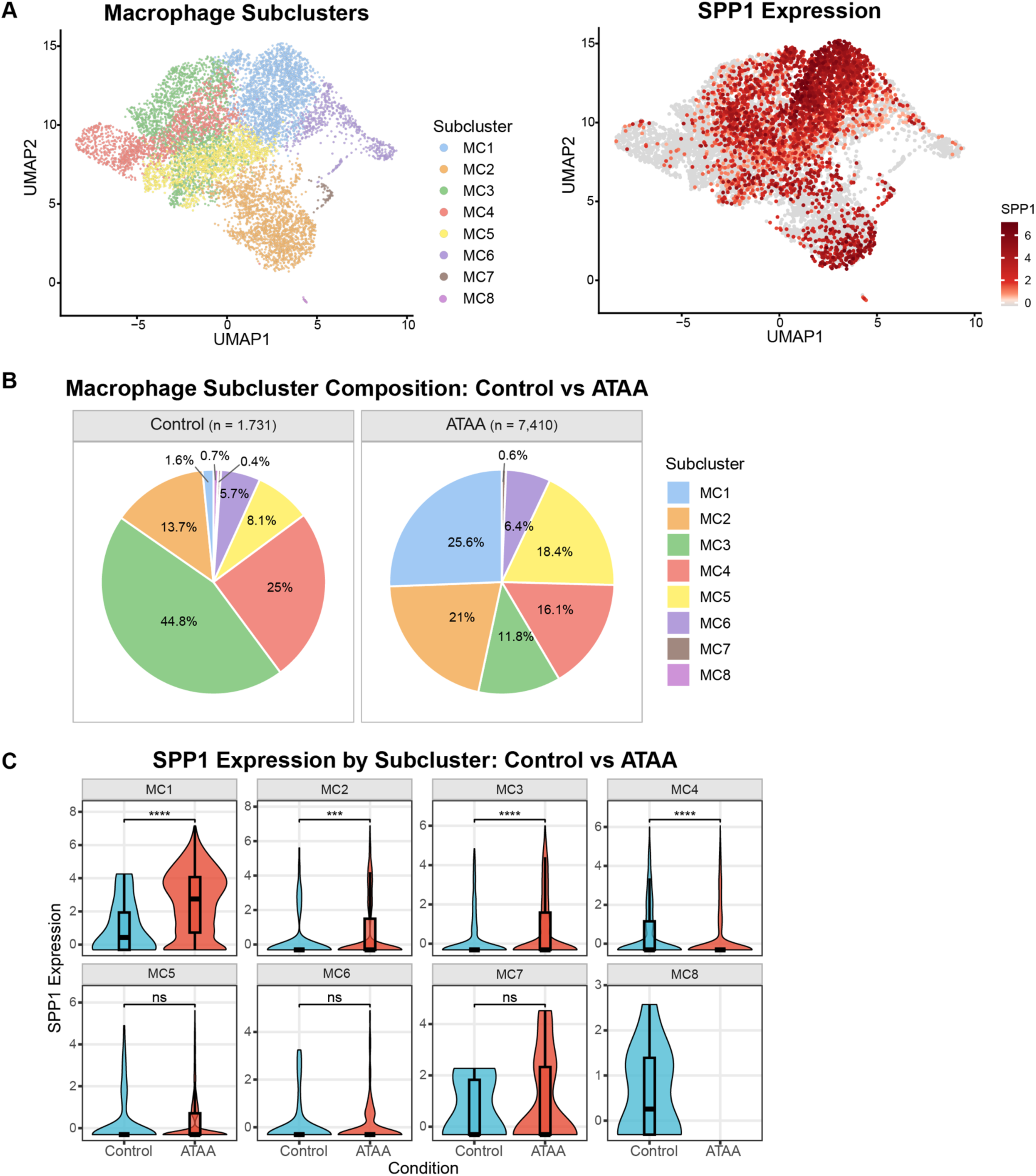
Identification of an osteopontin-high macrophage subpopulation in human ascending thoracic aortic aneurysm. A. UMAP visualization of 9,141 macrophages after subclustering. Left panel shows cells colored by subcluster identity (MC1–MC8); right panel shows *SPP1* expression levels with a continuous color scale from gray (low) to dark red (high). B. Pie charts showing the distribution of macrophage subclusters in Control (n = 1,731) and ATAA (n = 7,410) samples. In Control, MC3 (44.8%) and MC4 (25.0%) predominate, whereas in ATAA, MC1 (25.6%), MC2 (21.0%), and MC5 (18.4%) are the most prevalent subclusters. MC8 is absent in ATAA. C. Violin plots showing *SPP1* expression levels in each macrophage subcluster, comparing Control (blue) and ATAA (red). Statistical comparisons were performed using the Wilcoxon rank-sum test. MC1 and MC3 showed highly significant upregulation in ATAA (\*\*\*\**p* < 0.0001), and MC2 showed significant upregulation (\*\*\**p* < 0.001). In contrast, MC4 showed highly significant downregulation in ATAA (\*\*\*\**p* < 0.0001). MC5, MC6, and MC7 showed no significant difference (ns). MC8 lacked sufficient ATAA cells for comparison.

The composition of macrophage subclusters differed substantially between conditions (Figure 5B). In Control tissue, MC3 (44.8%) and MC4 (25.0%) together accounted for approximately 70% of all macrophages, whereas ATAA macrophages exhibited a markedly different distribution, with MC1 (25.6%), MC2 (21.0%), and MC5 (18.4%) representing the three largest subpopulations. Notably, MC1 was almost exclusively composed of ATAA-derived cells (1,898 of 1,925 cells; 98.6%), identifying it as an ATAA-specific macrophage subpopulation. Conversely, MC8 was composed entirely of Control-derived cells and was absent in ATAA (Control, n = 12; ATAA, n = 0).

Differential marker gene analysis and GO enrichment analysis revealed distinct functional programs across subclusters (Table 1, Supplementary Figure S6A & S6B). MC1, the ATAA-dominant subcluster with the highest *SPP1* expression, was characterized by marker genes *GAPDH*, *BCL2A1*, and *CD83*, and was enriched for stress response and immune system process pathways. The Control-dominant subclusters MC3 and MC4 were characterized by stimulus-responsive genes (*FKBP5*, *HERPUD1*, *CD163*) and phagocytic markers (*HLA-DRB5*, *TGFBI*, *CCL2*), respectively, and were enriched for stimulus response and endocytosis pathways. MC6, characterized by inflammatory markers *EREG*, *LYZ*, and *CXCL8* and enriched for defense response and lipopolysaccharide response pathways, represented an inflammatory macrophage population that expanded modestly in ATAA (Control, 5.7%; ATAA, 6.4%). MC8, the sole Control-specific subcluster, expressed tissue-remodeling markers *MFGE8*, *COL14A1*, and *SOD3* and was enriched for muscle system process pathways, suggesting a tissue-resident identity lost in disease.

**Table 1.**
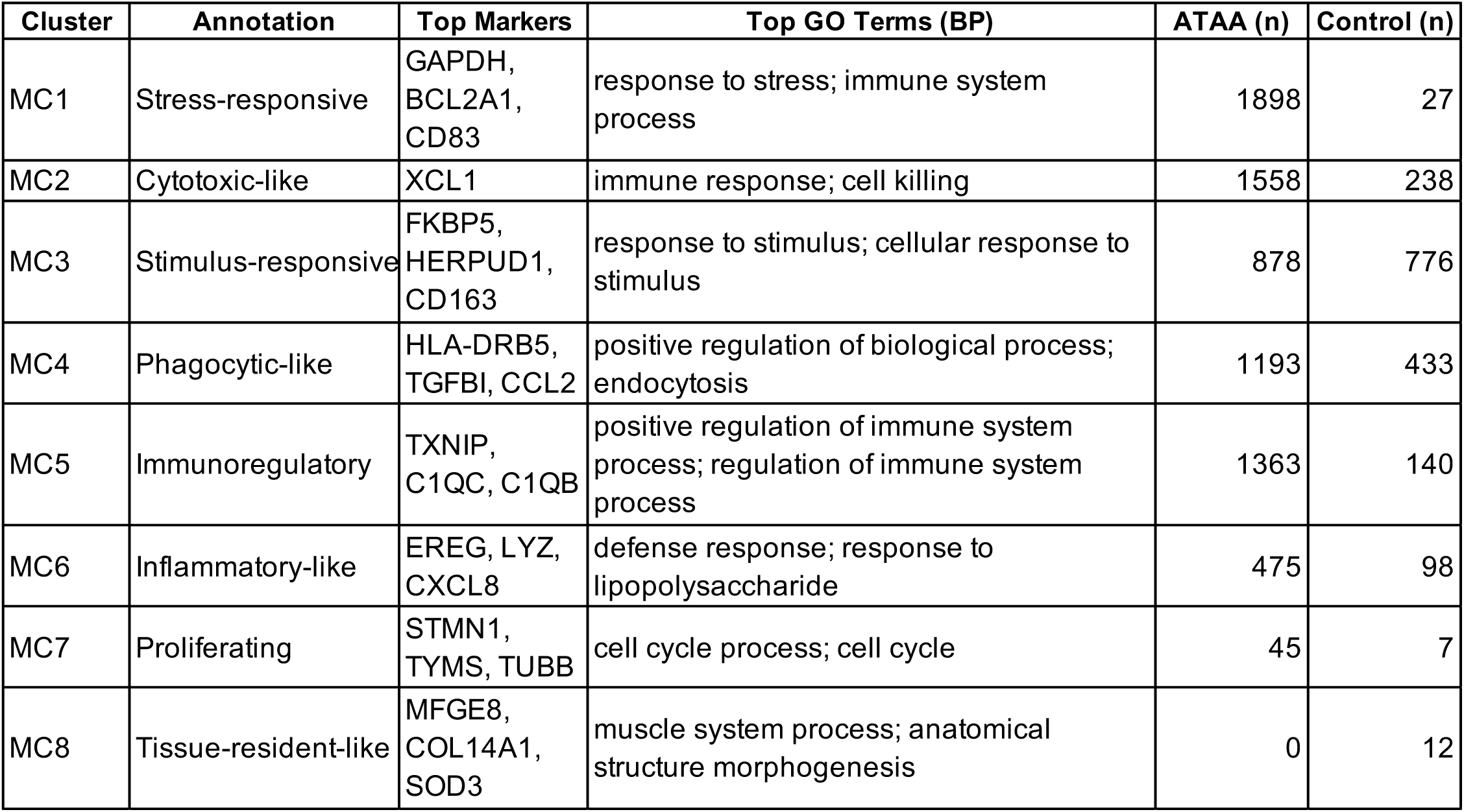
Summary of macrophage subcluster annotations, marker genes, enriched pathways, and cell composition in human thoracic aortic aneurysm. For each of the eight macrophage subclusters (MC1–MC8), the table lists the functional annotation, top marker genes ranked by differential expression score, top two significantly enriched Gene Ontology Biological Process (GO:BP) terms ranked by adjusted *p*-value, and the number of cells from ATAA and Control samples. Marker genes were selected from those with adjusted *p* < 0.05 and positive log₂ fold change. GO:BP terms were filtered for statistical significance (adjusted *p* < 0.05).

Subcluster-resolved analysis of *SPP1* expression revealed a heterogeneous pattern across macrophage subpopulations (Figure 5C). Significant *SPP1* upregulation in ATAA was observed in MC1, MC3 (both *p* < 0.0001), and MC2 (*p* < 0.001), while MC4 exhibited highly significant downregulation in ATAA (*p* < 0.0001). MC5, MC6, and MC7 showed no significant difference between conditions, and MC8 lacked sufficient ATAA-derived cells for statistical comparison. Collectively, these findings demonstrate that *SPP1* upregulation in ATAA macrophages is not a pan-macrophage response but is concentrated in MC1, an ATAA-expanded stress-responsive population, highlighting the subcluster-specific nature of OPN induction within the macrophage compartment.

## Discussion

In this study, we identified a defined pre-rupture period in AngII-treated *Fbn1*^mgR/mgR^ and characterized the molecular features of the aortic wall during this critical window. Single-section transcriptomic analysis revealed time-dependent gene expression changes preceding aortic rupture, among which osteopontin (*Spp1*/OPN) emerged as a notably upregulated gene at the pre-rupture timepoint. Immunofluorescence analysis confirmed markedly increased OPN protein expression in the aortic wall of *Fbn1*^mgR/mgR^ mice at 72 h after AngII infusion, while minimal expression was observed in WT mice or at earlier timepoints. Reanalysis of publicly available human ATAA single-cell RNA sequencing data further demonstrated that *SPP1* expression was elevated in aneurysmal tissue relative to non-aneurysmal controls, with macrophages exhibiting the most pronounced upregulation. Taken together, these findings highlight OPN as a molecular marker characterizing a pre-rupture state of the aortic wall that is conserved across mouse and human disease.

Prior studies on TAA have predominantly focused on mechanisms of aneurysm formation and progressive dilation, yielding important insights into extracellular matrix remodeling, smooth muscle cell dysfunction, and chronic inflammatory pathways^4,27^. In contrast, the molecular events immediately preceding aortic rupture remain poorly defined, largely because rupture occurs unpredictably and often catastrophically. The AngII-treated *Fbn1*^mgR/mgR^ mice model provides a unique opportunity to address this gap, as it exhibits a reproducible temporal course in which all animals survive to 24 h after AngII infusion, followed by progressive decline in survival. This reproducibility allowed us to define a pre-rupture window and analyze the aortic wall before overt rupture, complementing prior work on aneurysm formation by shifting attention toward the molecular transition from a vulnerable yet intact aortic wall to rupture.

Aortic rupture typically arises from localized structural failure rather than uniform deterioration of the entire vessel wall^28^. Previous transcriptomic and molecular studies of aortic disease have approached this tissue at different scales: bulk analysis of whole-tissue homogenates^5^, single-molecule spatial transcriptomics at single-cell resolution ^6^, and single-cell RNA sequencing^7,8^. While each approach has provided valuable insights, bulk analysis may dilute focal molecular signals by averaging across structurally heterogeneous regions, whereas single-cell and single-molecule spatial approaches, though highly resolved, profile a limited gene panel or lose tissue-level structural context. Single-section transcriptome analysis occupies an intermediate scale that preserves tissue-level structural information while enabling unbiased whole-transcriptome profiling of individual aortic wall sections, providing a novel framework for interrogating rupture-associated molecular alterations.

Cross-genotype heatmap analysis identified correlated gene clusters between WT and *Fbn1^mgR/mgR^*mice. Among these, WT C1 and *Fbn1*^mgR/mgR^ C2 showed concordant upregulation over time in both genotypes and were enriched for ciliary structure and mechanosensitive signaling ^29,30^, consistent with a shared hemodynamic response to AngII-induced wall stress. WT C3 and *Fbn1*^mgR/mgR^ C3 also showed correlation but with divergent trajectories: in WT mice, this cluster displayed a transient downregulation at 24 h followed by recovery at 72 h — consistent with a physiological homeostatic response — and was enriched for extracellular matrix organization and connective tissue development. In contrast, the corresponding *Fbn1^mgR/mgR^* cluster showed progressive downregulation without recovery approaching the pre-rupture timepoint, and was instead dominated by protein stabilization and proteostasis-related pathways, suggesting a progressive impairment of protein quality-control capacity in the aneurysmal aortic wall under acute hemodynamic stress.

OPN has previously been implicated in TAA pathology, primarily in the context of aneurysm formation and chronic inflammatory remodeling^31,32^. Notably, single-molecule spatial transcriptomics has recently identified a calcification-related smooth muscle cell subpopulation in human TAA^6^, and OPN is a well-established regulator of vascular calcification and osteochondrogenic transition^33^, raising the possibility that OPN upregulation in the pre-rupture aortic wall may reflect an osteochondrogenic shift within the smooth muscle cell compartment. Elevated *Spp1* expression in adventitial fibroblasts of TAA has also been reported, and circulating *SPP1* levels are elevated in TAA and abdominal aortic aneurysm patients^34–37^. However, whether OPN specifically marks the transition to an imminent rupture state — rather than reflecting aneurysm presence per se — remained unknown. The present study addresses this question directly by leveraging a model with predictable rupture timing, demonstrating that OPN upregulation occurs specifically at the pre-rupture timepoint (72 h) but not at the earlier timepoint (24 h). Given that OPN is a secreted protein, these findings raise the possibility that circulating OPN levels may specifically reflect rupture susceptibility rather than aneurysm presence alone, potentially contributing to a multi-parameter risk stratification framework.

Among the cell types showing *SPP1* upregulation in human ATAA, macrophages exhibited the most pronounced induction, suggesting that macrophage-driven *SPP1* — rather than the fibroblast-derived *SPP1* previously implicated in chronic aneurysm remodeling^36^ — may represent the dominant cellular source in the context of aortic wall stress and rupture susceptibility. Subcluster analysis of the macrophage compartment further revealed that this upregulation was not a pan-macrophage response but was concentrated in MC1, a stress-responsive subpopulation that expanded dramatically in ATAA (Control, 1.6%; ATAA, 25.6%) and was almost exclusively composed of ATAA-derived cells (98.6%). MC1 was characterized by marker genes *GAPDH, BCL2A1*, and CD83, and was enriched for stress response and immune system process pathways, suggesting that OPN induction is coupled to specific stress-activated programs in this subpopulation. Although *SPP1*-positive macrophage subpopulations have been identified in ATAA^32^, perivascular adipose tissue of heart failure patients^38^, and atherosclerotic plaques^39^, the present study provides the first characterization of a stress-responsive macrophage subpopulation — dramatically expanded in ATAA and almost exclusively composed of ATAA-derived cells — in the context of pre-rupture aortic wall stress.

This study has several limitations. The tissue-level transcriptomic approach used in the mouse model did not permit identification of the specific cellular sources of OPN within the aortic wall, and the functional role of OPN in rupture progression remains to be established through targeted experimental models. Furthermore, human single-cell RNA sequencing data were derived exclusively from non-ruptured ATAA surgical specimens, and the temporal proximity of these samples to potential rupture events could not be determined.

## Supporting information

Supplementary Figure S1. UpSet plot analysis of differentially expressed genes across pairwise comparisons.

## Acknowledgements

This work was supported by Grants-in-Aid for Scientific Research from the Japan Society for the Promotion of Science (JSPS KAKENHI Grant Numbers JP21K15366 and JP25K18723), Toyota Physical and Chemical Research Institute Scholar, Kowa Life Science Foundation, and Mochida Memorial Foundation for Medical and Pharmaceutical Research to K.S.; JSPS KAKENHI (Grant-in-Aid for JSPS Fellows, Grant Number JP24KJ2100) to Y.S.; and the Platform Project for Supporting Drug Discovery and Life Science Research (Basis for Supporting Innovative Drug Discovery and Life Science Research [BINDS]) from AMED (Grant Number JP21am0101104) to H.T.

