## Supplementary figures and images for "Osteopontin Upregulation Defines a Pre-Rupture State in Thoracic Aortic Aneurysms in Mice and Humans"

### Supplementary Figure S1. UpSet plot analysis of differentially expressed genes across pairwise comparisons.

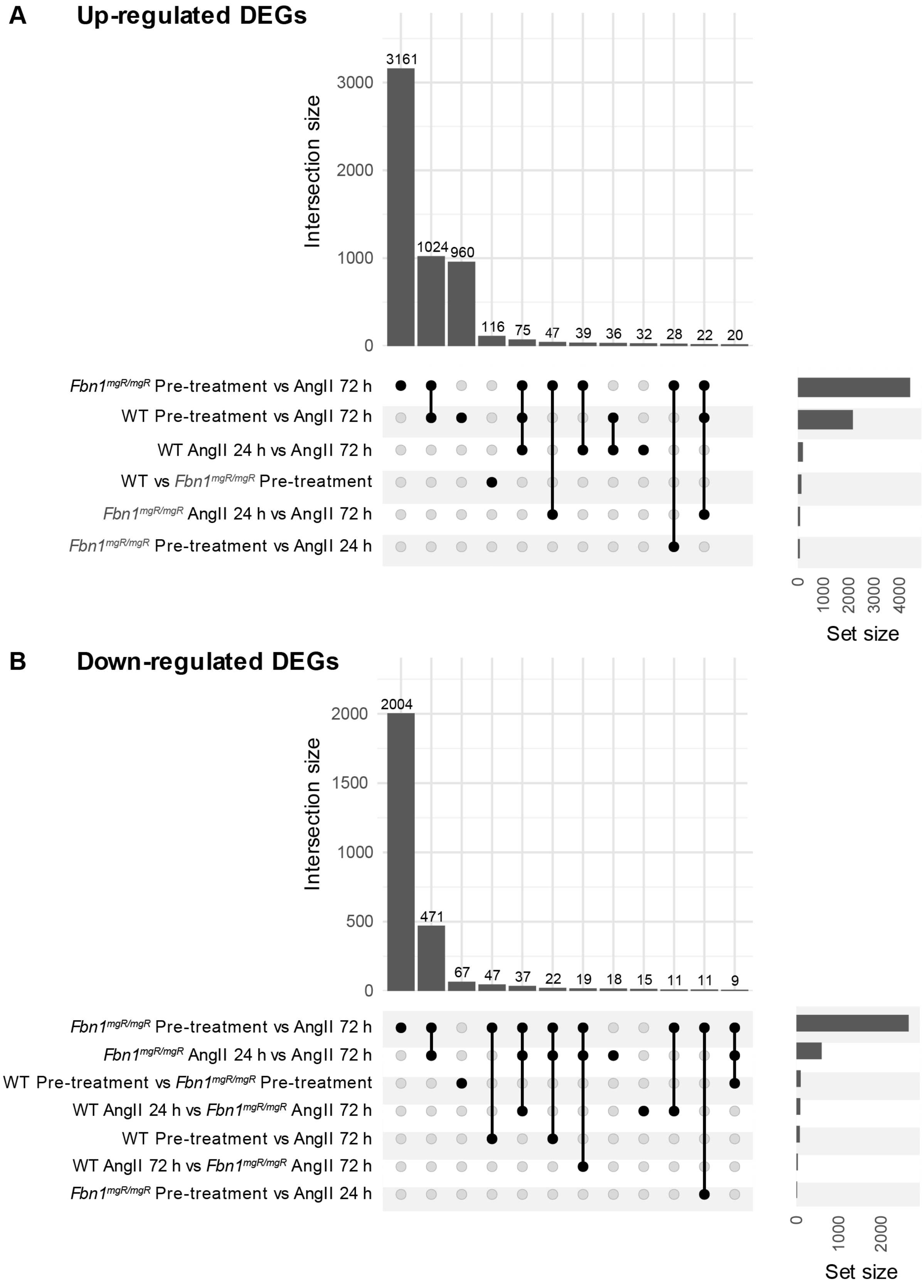
